# How Visual Experience Shapes Face Processing: Divergent Representational Strategies Emerge from Specialized and Diverse Visual Diets in Artificial Neural Networks

**DOI:** 10.64898/2026.02.01.703162

**Authors:** Xiqian Lu, Yi Jiang

**Affiliations:** State Key Laboratory of Cognitive Science and Mental Health, Institute of Psychology, Chinese Academy of Sciences, Beijing 100101, China; Department of Psychology, University of Chinese Academy of Sciences, Beijing 100049, China

## Abstract

Face perception in humans is supported by specialized neural mechanisms, yet the developmental origins of these mechanisms remain debated. We trained two artificial neural networks (ANNs) on identical orientation-classification tasks but with distinct visual diets: one exposed exclusively to adult faces (Face-ANN) and the other to diverse object categories (Object-ANN). Face-ANN acquired human-like face-processing signatures, including robust performance on Mooney faces, infant faces, and minimal three-dot patterns, and a reliance on low-frequency global structure. However, it failed to recognize face-like objects or most animal faces (except for monkeys), indicating limited overgeneralization. Object-ANN showed the opposite profile, excelling at face-like objects and animal faces but failing on abstract faces. These dissociations demonstrate that domain-specific visual experience alone can give rise to core properties of human face perception, while broad visual experience supports flexible generalization and face pareidolia. Our findings highlight how distinct visual diets shape representational strategies and offer a computational framework for probing the evolutionary origins of face-selective systems.

## Introduction

Human vision is not a passive reflection of the external world, but an active and constructive process that can extract rich structure from visual input within a single glance^1^. Among its many specialized functions, face perception stands out as a particularly striking example. Humans exhibit exceptional sensitivity to facial configuration, supported by dedicated neural mechanisms rather than a general object recognition system^2–7^. This specialization is classically demonstrated by the face inversion effect: inverting a face leads to a pronounced decline in recognition performance and reduced engagement of face-selective processing, rendering face perception more object-like^3,8–14^. Sensitivity to faces emerges early in development, with infants already preferring face-like patterns^15–18^. At the same time, this highly efficient system is also prone to characteristic errors. For example, in face pareidolia, observers readily perceive illusory faces in clouds, rocks, or coffee stains, highlighting both the power and the fallibility of face-selective visual mechanisms^19–22^.

What gives rise to this remarkable specificity in face processing? Although the longstanding debate over whether face-selective mechanisms are innate or acquired through development remains unresolved^23,24^, a more fundamental question concerns the nature of the experience that shapes face selectivity. Is face-specific processing primarily driven by extensive, domain-specific exposure to faces, or can it emerge from general visual experience shaped by the statistical structure of natural images^25–28^?

To address this question, we adopt convolutional neural networks as computational models of visual representation learning. These models provide precise control over visual input and implement hierarchical processing architectures that share important organizational principles with the primate visual system^29–38^. Artificial neural networks (ANNs) were trained with identical architectures and optimization objectives — a self-supervised orientation prediction task — while differing only in their visual input statistics. Compared with supervised learning that relies on explicit labels, self-supervised learning provides a closer approximation to how human visual experience is acquired. Specifically, we compared a face-enriched visual diet with a general natural image diet. This controlled computational framework allows us to isolate the contribution of visual input distributions to the emergence of face-selective representations, offering mechanistic insight into the origins of face processing specificity^39,40^.

## Results

We adopted the AlexNet architecture and modified the optimization task to classify image orientations (0°, 90°, 180°, and 270°). Although this four-way orientation classification task appears simple, it is sufficient to induce robust visual representations^41^. One model was trained exclusively on face images (Face-ANN), while the other was trained on a diverse set of general object categories (e.g., animals, vehicles, tools; Object-ANN; Fig. 1). Both models were then evaluated on the same stimulus set, allowing their performance to be directly compared.

**Fig 1.**
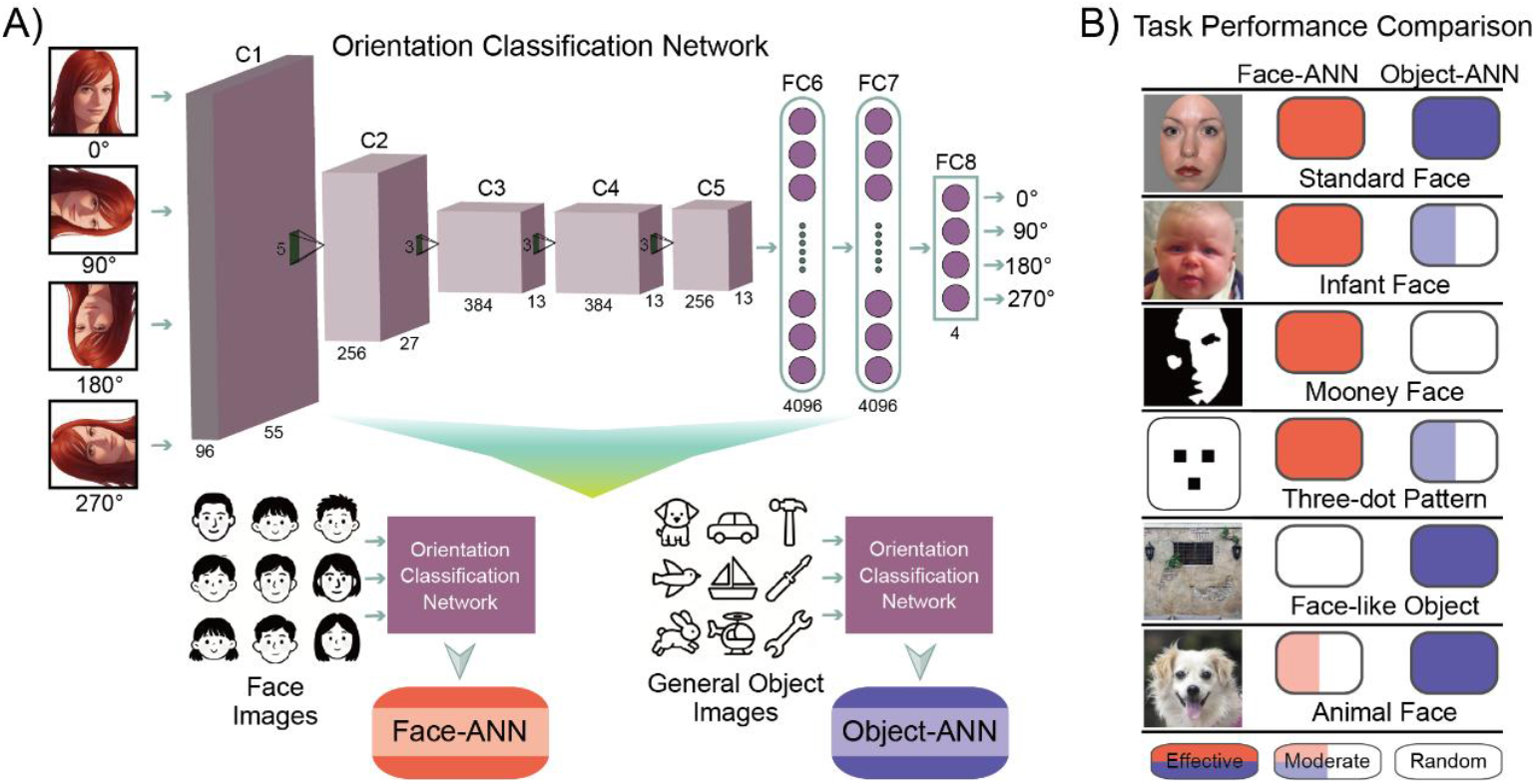
Overview of methods and key results. (A) Schematic of the model architecture and optimization task. We adopted an AlexNet architecture trained on a four-way orientation classification task (0°, 90°, 180°, and 270°). Two models were trained: *Face-ANN*, which received only face images, and *Object-ANN*, which received general object images. (B) Comparison of task performance. Face-ANN exhibits robust face-processing abilities, successfully recognizing faces even in highly abstract forms, such as Mooney faces and three-dot patterns, but fails to process face-like objects. In contrast, Object-ANN cannot recognize abstract faces, yet performs well on face-like objects and animal faces.

### Both Face-ANN and Object-ANN Can Process Faces, but They Rely on Distinct Representational Strategies

We first evaluated both models using standard face images. These faces were presented within an identical elliptical mask on a neutral gray background, containing no contextual or non-face cues. Thus, classification relied solely on facial information. Both Face-ANN and Object-ANN achieved perfect classification accuracy (1.00) for these images (Fig. 2A).

**Fig 2.**
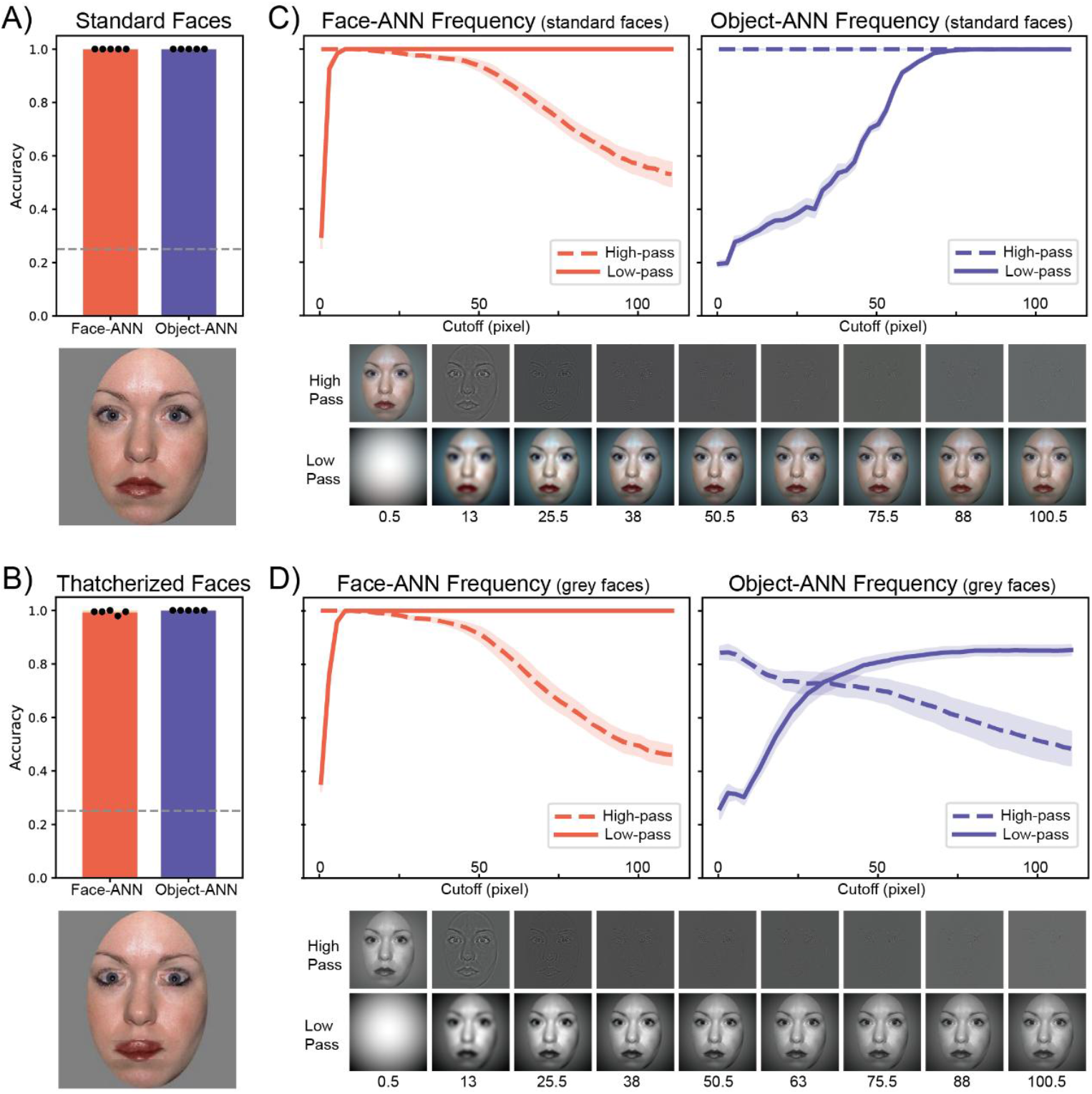
Model performance on standard faces. Both Face-ANN and Object-ANN show perfect performance on (A) standard faces and (B) Thatcherized faces. (C) The two models diverge in their sensitivity to spatial frequency: Face-ANN is robust to the loss of high-frequency information, whereas Object-ANN is robust to the loss of low-frequency information. (D) Spatial frequency effects persist when faces are converted to grayscale.

To further examine how the models represent facial structure, we generated face images with Thatcher illusion configuration by inverting the eyes and mouth of standard faces. This manipulation preserved the global configuration while disrupting local features. Both Face-ANN (0.99) and Object-ANN (1.00) classified these Thatcherized faces perfect performance (Fig. 2B), suggesting that both rely primarily on global facial configuration rather than local feature orientation.

The two models diverged sharply, however, in their dependence on spatial frequency content. We tested the models with low-pass and high-pass filtered versions of standard faces (Fig. 2C). Across different cutoff frequencies, Face-ANN remained highly robust when high-frequency information was removed but was impaired by the loss of low-frequency components. In contrast, Object-ANN’s performance remained stable when low-frequency information was removed but declined markedly when high-frequency components were eliminated. Considering that spatial frequency is closely tied to luminance variation, we also tested grayscale versions of the original RGB images (Fig. 2D). Grayscale conversion barely affected Face-ANN’s overall accuracy or its performance across spatial frequencies. However, Object-ANN showed a slight reduction in overall accuracy and a pronounced drop under low-frequency removal conditions. Together, these results suggest that although both models can recognize faces, they rely on fundamentally different visual representations. Face-ANN depends more on low-frequency, holistic facial structure, whereas Object-ANN is more sensitive to high-frequency content.

### Face-ANN Processes Mooney Faces in a Human-like Manner

Mooney faces—highly abstract black-and-white stimuli containing only coarse, low-frequency information—are readily recognized by humans^42^. We tested both models with Mooney faces (including frontal and profile views) and their matched non-face controls (Fig. 3A)^43,44^. Face-ANN classified the orientation of Mooney faces with high accuracy (0.80) but performed poorly on the non-face controls (0.44), despite their similar patch composition. In contrast, Object-ANN failed to classify either Mooney faces (0.26; *Z* = 2.64, *p* = 0.008) or controls (0.17; *Z* = 2.61, *p* = 0.009) above chance.

**Fig 3.**
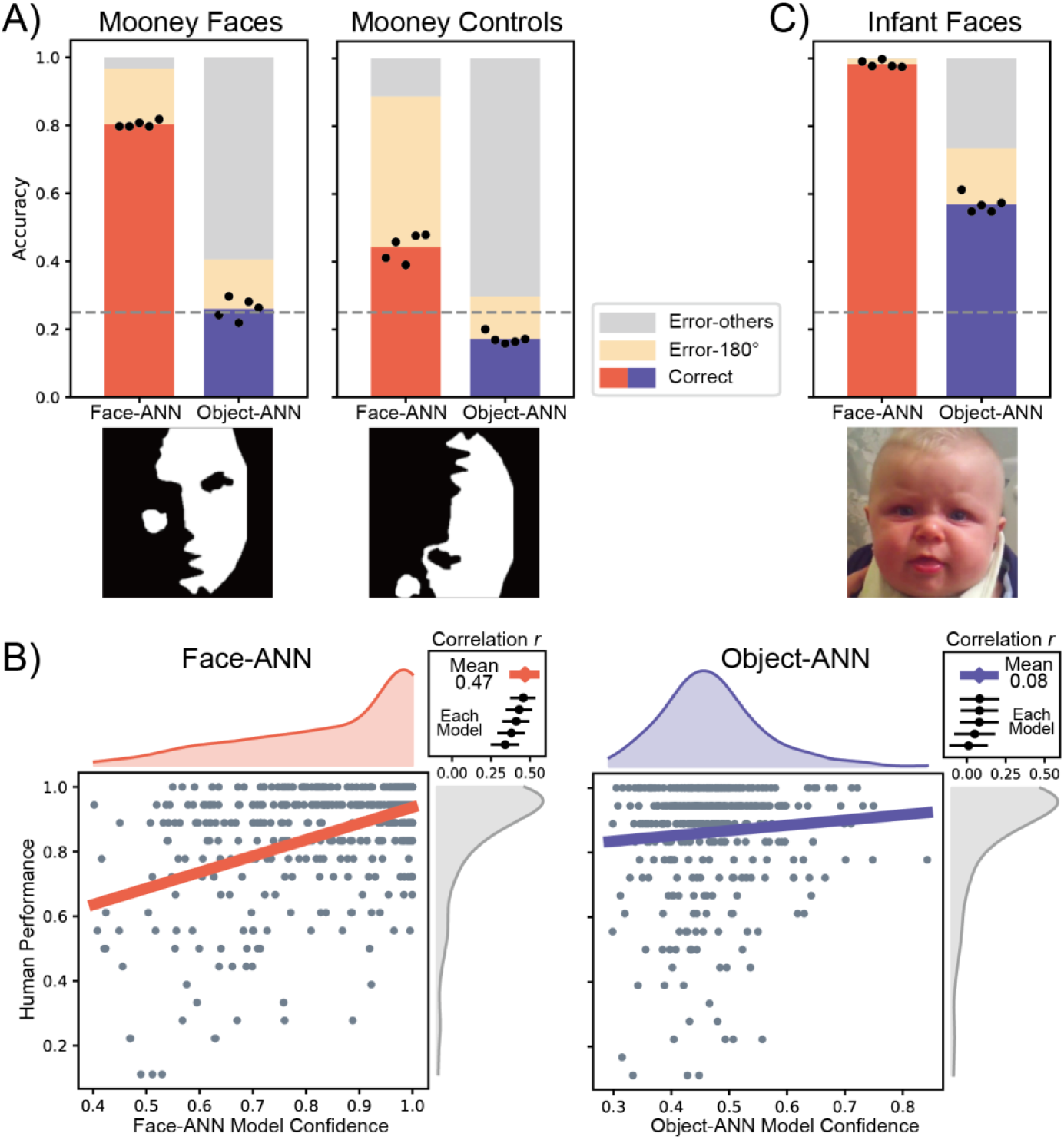
Model performance on Mooney and infant faces. (A) Model performance on Mooney faces and their matched non-face controls. Black dots represent results from individual models trained with different random seeds. Error types are separated into misclassifications as the 180°-rotated version and all other errors. (B) Correlation between model performance and human perception for Mooney faces. Marginal histograms show the distributions along each axis. Correlations are shown for both the mean model performance and individual models trained with different random seeds (top-right inset). (C) Model performance on infant faces.

We further compared model performance with human behavioral data (Fig. 3B). The model index was defined as the confidence for correctly classified images, and the human index was the proportion of correctly judged upright Mooney faces. Face-ANN’s confidence scores correlated significantly with human responses (r = 0.47, p < 0.001), whereas Object-ANN showed no such relationship (r = 0.08, *p* = 0.11; Fisher z-test: *Z* = 6.34, *p* < 0.001). These findings suggest that Face-ANN processes Mooney faces in a manner closely aligned with human perception.

### Face-ANN Generalizes to Infant Faces Despite Training Exclusively on Adult Faces

We tested both models with infant face images (Fig. 3C). Infant faces differ markedly from adult faces, exhibiting a distinct “baby schema” characterized by a disproportionately large forehead, rounder eyes positioned lower on the face, smaller nose and mouth, and fuller cheeks. Although Face-ANN was trained exclusively on adult faces, it classified infant faces with near-perfect accuracy (0.98). Object-ANN could also process infant faces but with significantly lower accuracy (0.57; *Z* = 2.63, *p* = 0.009). These results indicate that the facial representations learned by Face-ANN from adult faces generalize robustly to infant faces.

### Face-ANN Interprets Minimal Three-dot Patterns as Faces

Next, we tested both models using very simple three-dot patterns, which consist of three squares arranged to mimic the spatial configuration of eyes and mouth. Such patterns have been shown to elicit face perception in both human infants^15^ and newborn chicks^45^. We created two contrast-polarity versions of these patterns that were geometrically identical, one set with light background and dark features, and another with dark background and light features. Natural faces typically have dark features on a light background.

We first tested three-dot patterns where the features were placed at appropriate locations but varied in size and introduced with positional jitter (Fig. 4A and 4B). Face-ANN classified light-background patterns well (0.80) but performed worse on dark-background patterns (0.34). Object-ANN showed moderate performance on both light-background (0.58; *Z* = 2.61, *p* = 0.009) and dark-background patterns (0.42; *Z* = 1.57, *p* = 0.12).

**Fig 4.**
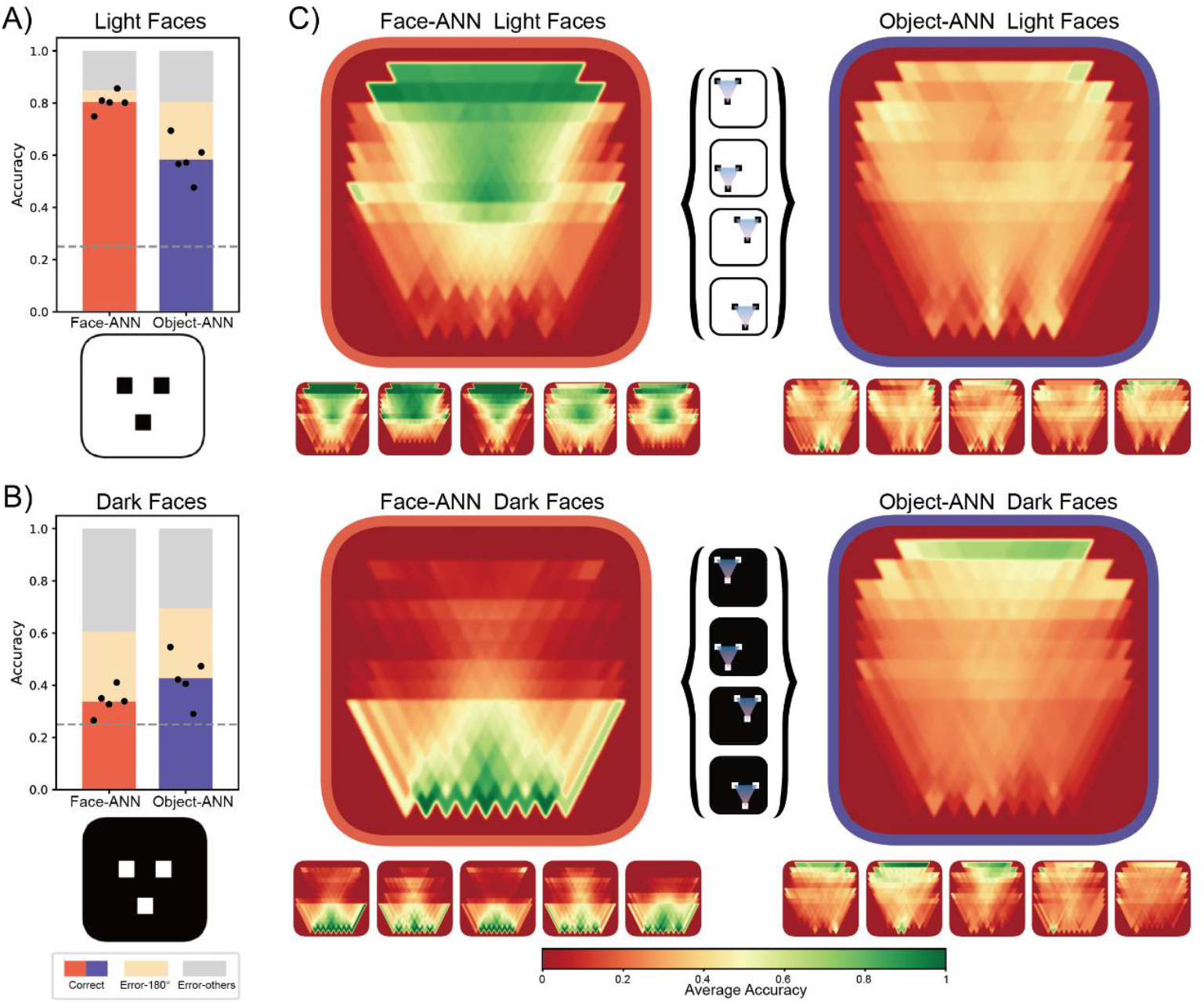
Model performance on three-dot patterns. The models show better performance on (A) light-background than on (B) dark-background three-dot face-like patterns. (C) Cumulative accuracy heatmaps for each model across light and dark patterns. Face-ANN exhibits distinct and contrast-dependent response profiles, whereas Object-ANN is not reliably modulated by contrast polarity. Small heatmaps depict results from individual models trained with different random seeds.

To examine spatial constraints, we systematically varied the absolute positions of the three-dot patterns while preserving their triangular configuration. Cumulative accuracy heatmaps (Fig. 4C) revealed that Face-ANN’s performance on light-background patterns peaked when features were located in the upper visual field, whereas performance on dark-background patterns peaked in the lower field. In contrast, Object-ANN’s accuracy was not reliably modulated by either feature location or contrast polarity. These findings suggest that Face-ANN, but not Object-ANN, interprets these minimal stimuli as faces and is sensitive to their natural contrast polarity.

### Object-ANN outperforms Face-ANN on Artificial Face-like Paintings

We next tested Arcimboldo paintings, which were created by a 16th-century Italian artist Giuseppe Arcimboldo. These impressive portrait paintings depict human faces composed entirely of non-face objects such as fruits, animals, and flowers. These compositions retain the overall structure of a face while possessing entirely non-face textures. Some of these works are reversible: they appear as faces when upright but transform into still life scenes (e.g., a bowl of vegetables) when inverted. We tested both models on these paintings (Fig. 5A). Object-ANN accurately classified the orientations of all paintings (1.00), including the reversible ones. Face-ANN also classified many of these paintings significantly above chance (0.57; *Z* = 2.81, *p* = 0.005). These results indicate that Face-ANN can partially generalize its facial representations to artificial compositions that preserve global facial structure.

**Fig 5.**
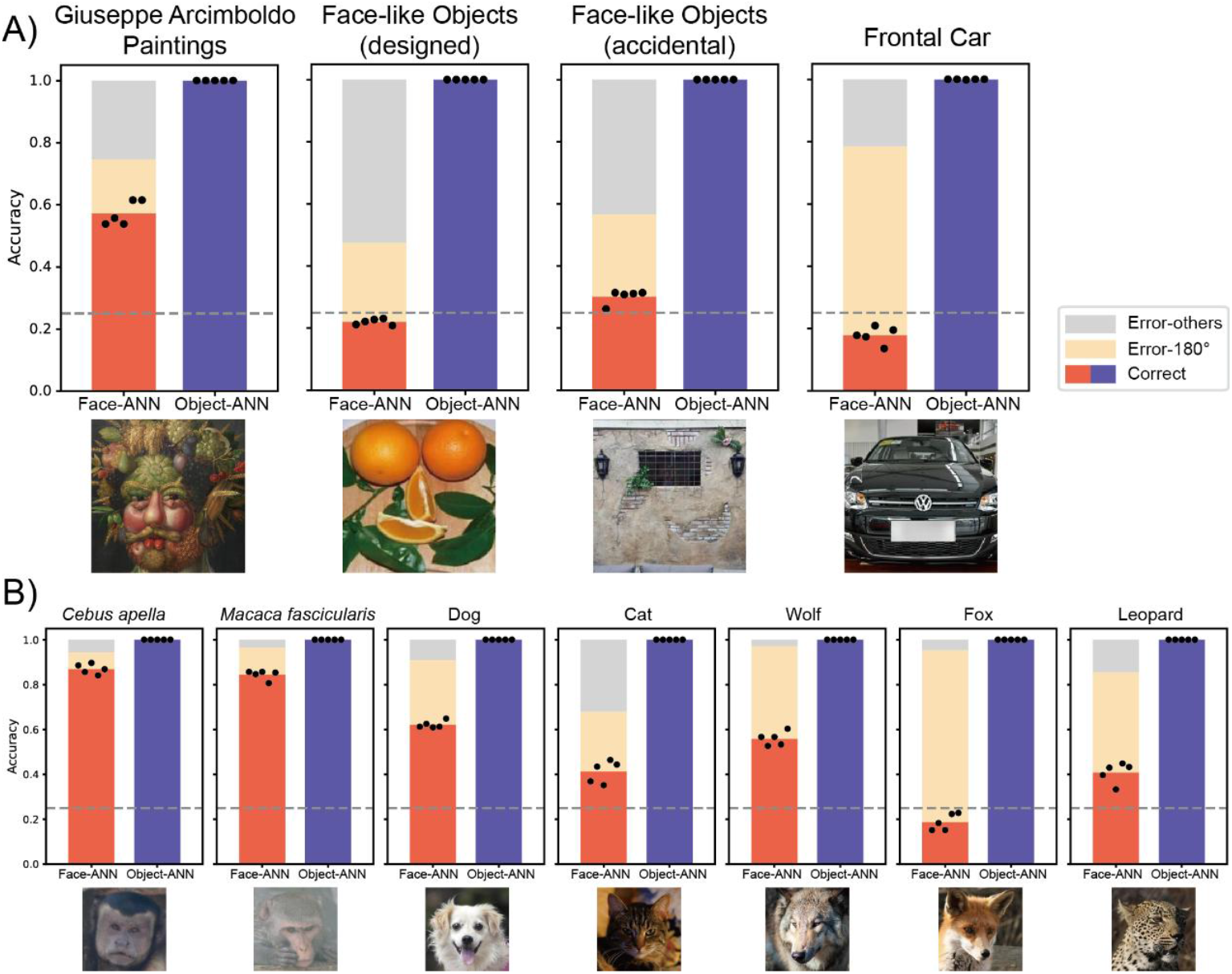
Model performance on face-like objects and animal faces. (A) Object-ANN achieves perfect accuracy on both designed and accidental face-like objects, whereas Face-ANN performs well only on Arcimboldo paintings. (B) Object-ANN also classifies all animal faces perfectly, while Face-ANN shows high accuracy only for monkey faces.

### Object-ANN, but not Face-ANN, Supports Face Pareidolia

Humans often experience face pareidolia—the perception of faces in inanimate objects. To examine whether the models exhibit similar tendencies, we tested their performance on two categories of face-like objects (Fig 5A). The first category (accidental) comprised naturally occurring objects that resemble faces (e.g., a house with two circular windows and a door). The second category (intentionally designed) consisted of artificial arrangements of objects (e.g., food items) deliberately organized to form face-like configurations. Face-ANN failed to classify either type of face-like object (designed: 0.22; accidental: 0.30). In contrast, Object-ANN correctly classified both types of face-like objects with perfect accuracy (designed: 1.00, *Z* = 2.79, *p* = 0.005; accidental: 1.00, *Z* = 2.80, *p* = 0.005).

We also evaluated frontal car images (Fig. 5A), which contain a regular visual pattern (headlights and a windshield arranged in a stable configuration) that may exhibit a statistical regularity analogous to facial structure. Object-ANN again achieved perfect classification accuracy (1.0), while Face-ANN performed below chance level (0.18; *Z* = 2.70, *p* = 0.007), frequently misclassifying cars as vertically oriented. These results suggest that Face-ANN does not overgeneralize its facial representations to either designed or accidental face-like objects, whereas Object-ANN might exploit subtle face-like structure in complex natural scenes, potentially enabling face pareidolia.

### Object-ANN Processes Animal Faces, Whereas Face-ANN Generalizes Only to Monkeys

Finally, we tested both models on a range of animal faces (Fig. 5B), including domestic species (dogs and cats), related wild species (foxes, wolves, leopards), and two primate species (*Cebus apella and Macaca fascicularis*). Object-ANN correctly classified all animal faces (1.00). Face-ANN, however, performed poorly on most animal species, showing low accuracy for cats (0.41; *Z* = 2.79, *p* = 0.005), foxes (0.19; *Z* = 2.80, *p* = 0.005), and leopards (0.41; *Z* = 2.79, *p* = 0.005), and relatively better performance for dogs (0.62; *Z* = 2.80, *p* = 0.005) and wolves (0.56; *Z* = 2.80, *p* = 0.005). Notably, Face-ANN performed well on monkey faces (*Cebus apella*: 0.89, *Z* = 2.79, *p* = 0.005; *Macaca fascicularis*: 0.84, *Z* = 2.80, *p* = 0.005). Across species, Face-ANN tended to misclassify images as vertically oriented. These results indicate that although human and animal faces share broad structural similarities, visual experience with human faces does not generalize widely to other species, except for closely related primates. Object-ANN, by contrast, benefits from its diverse visual diet and can process animal faces robustly.

## Discussion

We evaluated two ANNs across a broad set of stimuli that humans intuitively categorize as faces. The results revealed a striking dissociation between the two models. Face-ANN exhibited remarkably human-like responses to abstract face stimuli. It successfully processed highly abstract Mooney faces and minimal three-dot face-like patterns, with performance closely paralleling human perception. However, it showed no evidence of face pareidolia and only limited sensitivity to animal faces. In contrast, Object-ANN, despite exhibiting excellent performance across diverse visual categories, failed to process abstract face stimuli altogether.

Face-ANN appears to have acquired an abstract, structural representation of faces through extensive orientation learning on face images, enabling strong generalization within the face domain. Notably, it achieved near-perfect performance on infant faces, which it had never encountered during training, and showed robust sensitivity to highly abstract face stimuli. Crucially, this face-specific processing did not rely on high-frequency facial details. Instead, the model learned invariant facial structure from face-only input, spontaneously emphasizing global configuration and core facial features. Consistent with this, Face-ANN depended minimally on high spatial frequency information and operated effectively using only low spatial frequency cues. Converting images from RGB to grayscale did not impair performance, indicating that Face-ANN primarily relies on luminance-based spatial structure rather than color information. This property was most clearly demonstrated in the Mooney face task. Mooney faces convey facial structure almost exclusively through coarse, low-frequency information. Face-ANN not only processed Mooney faces successfully, but its performance was also significantly correlated with human perception. Although the model could exploit high spatial frequency information when low-frequency cues were absent, this capacity was comparatively limited.

A similar human-like sensitivity emerged for another type of abstract stimuli: T-shaped three-dot face configurations. Face-ANN exhibited distinct processing of geometrically identical configurations that differed solely in contrast polarity, resulting in markedly different performance for light-background versus dark-background patterns. Specifically, faces presented against a light background with darker local features—mimicking the natural luminance configuration of real human faces—elicited superior performance, especially when facial features were concentrated in the upper visual field. When the configuration was disrupted and only isolated “eyes” or a “mouth” were retained, performance dropped sharply (see Fig. S1). This pattern closely mirrors well-documented human perceptual biases in early face processing^16,46^.

Despite these strengths, Face-ANN’s face processing abilities did not generalize to objects that humans subjectively perceive as face-like. Face-ANN maintained relatively high performance on face-like paintings, which preserve facial structure despite differences in material properties. However, it failed to process face-like objects in natural scenes, even when these objects contained a T-shaped configuration. This failure occurred despite the geometric similarity between these objects and the three-dot patterns, indicating that Face-ANN does not overgeneralize its facial representations and spontaneously extract face-like structure from complex visual scenes. Interestingly, Face-ANN consistently misclassified upright cars as inverted stimuli, possibly because the lower placement of headlights was misinterpreted as upper facial features of an inverted face. These findings suggest that pure facial experience produces a highly precise feature detector rather than a hallucination-prone module. Human face-processing phenomena such as pareidolia may therefore not arise from face-specific experience alone, but instead reflect interactions among multiple visual systems, for example, between rapid, coarse detection mechanisms and fine-grained recognition processes. This interpretation aligns with two recent findings: first, mainstream face detection models do not exhibit face pareidolia, whereas fine-tuning animal face detection models more readily induces pareidolic responses^47^; second, ANNs that included object categorization in their training tasks represented pareidolia faces more similarly to human neural responses than those that did not^26^.

Object-ANN, by contrast, benefited from broad visual experience and accordingly showed superior performance across a wider range of image categories. Although it was never explicitly trained on faces, it processed standard faces perfectly and was unaffected by local feature inversion. It also showed excellent performance on animal faces and face-like objects. Importantly, we tested not only naturally occurring face-like objects but also artificially constructed face-like configurations that, in principle, lack an intrinsic orientation unless interpreted as faces. Nevertheless, these stimuli were also processed perfectly by Object-ANN. One possible explanation is that these images were deliberately photographed from canonical viewpoints and contained subtle directional cues, such as shading or lighting gradients, that Object-ANN could exploit. This interpretation may also account for why Object-ANN performed well on paintings. Although the elements in paintings are arranged subjectively by artists, they often retain implicit orientation cues, such as oblique light sources or gravity-consistent object placement. Because Object-ANN relies heavily on high spatial frequency information, it may be particularly sensitive to such local visual cues, while remaining ineffective when confronted with abstract face stimuli that lack high-frequency detail.

Together, our findings offer a new perspective on a longstanding question in face perception research. Human face processing is widely understood as a complex system composed of multiple components, with some elements shaped by postnatal experience and others reflecting evolutionary specialization. Even the so-called innate components can be viewed as “experience-based,” reflecting millions of years of evolutionary history and instantiated in specialized neural substrates such as the fusiform face area (FFA). By training ANNs to approximate specific forms of visual experience, we show that face-specific experience alone can give rise to hallmark properties of human face processing, including low-frequency dominance, sensitivity to Mooney faces, and face-like interpretation of minimal configurations such as three-dot patterns. These results highlight the potential of ANNs as a powerful tool for probing longstanding questions about the evolutionary origins of cognitive systems—questions that are often difficult to address directly using traditional empirical approaches. More broadly, they illustrate how evolutionary environments may shape the fundamental mechanisms underlying human perceptual experience.

## Method

### Model Training

We implemented AlexNet^48^ in Python (3.12.11) using PyTorch (2.8.0) and modified the final classification layer to output four orientation classes. Instead of using the original labels, we generated four copies of each training image by rotating it to four orientations (0°, 90°, 180°, and 270°), and used these orientations as the training labels. Image rotations were implemented using combinations of transpose and flip operations^41^: for a 90° rotation, we first transposed the image and then flipped it vertically; for a 180° rotation, we first flipped the image vertically and then horizontally; and for a 270° rotation, we first flipped the image vertically and then transposed it. All training images were preprocessed to 224×224 pixels. For images larger than this size, we resized the shorter side to 224 pixels and cropped the center. For smaller images, we centered them on a 224×224 black background.

We trained two models: one using the VGGFace2 training set^49^, referred to as *Face-ANN*; and the other using the ImageNet Large Scale Visual Recognition Challenge (ILSVRC)-2017 training set^50,51^, referred to as *Object-ANN*. (Control analyses addressing differences in dataset size between the two datasets are reported in Table S1). Each model was trained from five different random initial seeds, and all reported results are the averages across the five models. Two of the five models were initially trained for 90 epochs. After observing that all model performances peaked before 40 epochs, the remaining three models were trained for 40 epochs. Training employed stochastic gradient descent (SGD) with momentum 0.9, weight decay 5e-4, and an initial learning rate of 0.01 that decayed according to a step learning rate schedule (step size = 10, gamma = 0.1). The batch size was set to 128. However, since all four rotated versions of each image were fed into the network simultaneously, each training batch effectively contained 512 images^41^. For comparison, we also employed untrained AlexNet models initialized with the same five random seeds. Since nearly all untrained models classified every test image as a single orientation, we have omitted these detailed results from the paper, but they are available in the raw data.

### Test Stimuli

#### Standard faces

were selected from the NimStim dataset^52^. We used only neutral and calm facial expressions, totaling 159 images. The original faces were presented on a neutral grey background and cropped within an elliptical mask to reveal only the face. Each image was resized so that its longer side was 224 pixels and then centered on a 224×224 neutral gray background.

#### Thatcherized Faces

were generated from the standard faces by inverting the eyes and mouth regions to create “Thatcher illusion”-like stimuli. Specifically, facial landmarks were detected using MediaPipe FaceMesh (https://ai.google.dev/edge/mediapipe/solutions/vision/face_landmarker) to localize the eye and mouth regions. The bounding boxes for the left eye, right eye, and mouth were computed based on their respective landmark points with appropriate padding, and each region was vertically flipped. All generated images were manually verified, with bounding box parameters fine-tuned when necessary to ensure precise alignment with the facial features while minimizing the inclusion of extraneous areas.

#### Infant faces

were selected from the Infant Annotated Faces (InfAnFace) dataset^53^. We first included only images labeled as non-rotated, non-occluded, non-expressive, and non-tilted. Each image was then manually cropped to a square facial region to ensure that only an upright infant face was shown, with minimal background and no other humans visible, and that the infant’s eyes were open. The cropped images were resized to 224×224 pixels, yielding a final set of 109 infant face images.

#### Mooney face

were obtained from the dataset by Schwiedrzik et al^43^. We used 96 upright Mooney faces and their corresponding scrambled control stimuli as originally designed. Additionally, we tested all 505 Mooney faces to compare the models’ responses with the human behavioral results reported in the original study. We compared the models’ confidence on correctly classified Mooney faces with the proportion of human observers who judged the upright face as upright. Each image was resized by scaling its longer side to 224 pixels and then centered on a 224×224 black background.

#### Face-like objects

were selected from “Faces in Things” dataset^47^. We first filtered images based on the dataset labels to include those containing only one easily recognizable face-like configuration, without emotional or amusing elements. We manually retained only images resembling a frontal human face (two eyes and one mouth) and excluded those resembling animal faces. Face regions were cropped according to the provided position labels; if the labeled region was non-square, the shorter side was expanded to form a square. We then manually excluded the images containing text. The images were then resized to 224×224 pixels. Based on the dataset labels with minor manual adjustments, we categorized the stimuli into two groups: accidental faces, where face-like patterns emerged naturally (e.g., a house with two circular windows and a door), and designed faces, where objects were intentionally arranged to resemble faces (e.g., food arranged to form facial features). This yielded 86 accidental and 78 designed face-like images.

#### Car

images were selected from the Comprehensive Cars dataset^54^. We first identified frontal-view images based on the dataset labels and then manually selected 160 images that contained no humans or text in the background. Each selected image was resized by scaling its shorter side to 224 pixels and then center-cropped to produce a 224×224 image.

#### Giuseppe Arcimboldo’s paintings

were obtained from Wikimedia Commons. We selected a subset of his works that depict faces composed of objects. The selected paintings are *Air, Autumn, Earth, Fire, Four Seasons in One Head, Rudolf II of Habsburg as Vertumnus, Spring, Summer, The Jurist, The Librarian, Water*, and *Winter*. For each painting, the facial region was manually cropped into a square, excluding the shoulders, and subsequently resized to 224×224 pixels. In addition, we included three reversible paintings, which exhibit a face-like appearance when oriented upright but transform into depictions of objects (such as a basin of vegetables) when inverted. These are *The Gardener, The Cook, and Fruit Basket*. Each of these was processed by scaling its longer side to 224 pixels and then centered on a 224× 224 black background.

#### Animal faces

stimuli were selected from two datasets. Dog, cat, fox, leopard, and wolf imaged were taken from the AFHQ dataset^55^, with 160 images for the first four species and 75 of wolves. Monkey images of Cebus apella and Macaca fascicularis were selected from the AFD dataset^56^. We manually excluded monkey images with pronounced head tilts that could obscure orientation or those showing food near the mouth. The final set included 162 Cebus apella and 75 Macaca fascicularis images. All images were originally square and resized to 224×224 pixels.

The filenames of all original images used in this study are listed in the supplementary materials.

### Orientation Classification Testing

Each test image was rotated to four orientations (0°, 90°, 180°, and 270°) and fed into each model, with the predicted label and associated confidence score were recorded for analysis. Mann-Whitney U test was used for model comparisons. (See Table S2 for grayscale-RGB comparisons of all stimuli tested in this study).

### Spatial Frequency Testing

For standard and Thatcherized faces, we test the influence of spatial frequency. Spatial frequency content was manipulated using a second-order Butterworth filter^57^. The filtering was implemented in the frequency domain via fast Fourier transform (FFT). Prior to filtering, images were padded using reflection and multiplied by a Hann window to minimize edge artifacts, which temporarily doubled their size. We applied cutoff frequencies ranging from 1 to 226 pixels in steps of 5, generating both low-pass and high-pass filtered versions for each cutoff. After filtering, images were cropped back to their original size. Each filtered image was then created four rotated versions (0°, 90°, 180°, and 270°), which were subsequently evaluated with the models.

### Three-dot Face Heatmap

Three-dot face stimuli were generated on a 224 ×224 white background. A rounded rectangle occupying 80% length served as the face outline. Eyes and mouth were represented as simple geometric squares. The mouth was positioned at a random height along the horizontal midline, while the eyes were symmetrically positioned above it at the same vertical height. Overlaps between the eyes and mouth were not allowed. A total of 300 valid configurations were generated. Each configuration was produced in two contrast versions, one with a white face and black features, and another reversely with black face and white features. The two types were structurally identical, differing only in color contrast, resulting in a total of 600 test images. These were input into each model, and the resulting orientation classification outputs were used to generate response heatmaps.

## Supporting information

Supplementary materials for the manuscript

## Acknowledgments

This study was supported by grants from the Brain Science and Brain-like Intelligence Technology – National Science and Technology Major Project (No. 2021ZD0203800), the National Natural Science Foundation of China (No. 32430043, 32500944), the Key Research and Development Program of Guangdong, China (2023B0303010004), and the Fundamental Research Funds for the Central Universities.

## Data availability

The trained model files, raw experimental data, and the list of stimulus images are available at https://doi.org/10.57760/sciencedb.psych.00977.

## Author Contributions

Y. J. and X. L. conceived and designed the study. X. L. performed model training, testing, statistical analysis, and wrote the original draft. All authors reviewed, edited, and approved the final manuscript for submission.

## Declaration of generative AI and AI-assisted technologies

During the preparation of this manuscript, the authors used DeepSeek and ChatGPT to assist with language editing and to enhance the overall readability of the text. All content generated with the assistance of this tool was thoroughly reviewed and revised by the authors, who take full responsibility for the final version of the manuscript.

