## Supplementary materials for the manuscript for "How Visual Experience Shapes Face Processing: Divergent Representational Strategies Emerge from Specialized and Diverse Visual Diets in Artificial Neural Networks"

Xiqian Lu[a,b], Yi Jiang[a,b]*

[a] State Key Laboratory of Cognitive Science and Mental Health, Institute of Psychology, Chinese Academy of Sciences, Beijing 100101, China

[b] Department of Psychology, University of Chinese Academy of Sciences, Beijing 100049, China

* Corresponding authors: Yi Jiang

**Supplementary Three-dot Patterns Test**

We tested modified versions of the original three-dot patterns, in which either the "eyes" or the "mouth" was removed. Compared with the intact three-dot patterns, both eye-removed and mouth-removed stimuli severely disrupted processing in the Face-ANN for light-background faces. In contrast, for the Object-ANN, only removal of the eyes led to a substantial performance impairment. Results for dark-background faces were more heterogeneous. Removing the mouth caused the Face-ANN to systematically misclassify the stimuli as the 180° inverted orientation, whereas the Object-ANN showed an increase in classification accuracy. By contrast, removing the eyes resulted in inconsistent outcomes across the five Face-ANN copies and reduced Object-ANN accuracy to nearly zero. Overall, for these minimal three-dot patterns, disrupting the core facial structure led to dramatic changes in model results, suggesting that models processing of the patterns is closely tied to their face-like structural configuration.

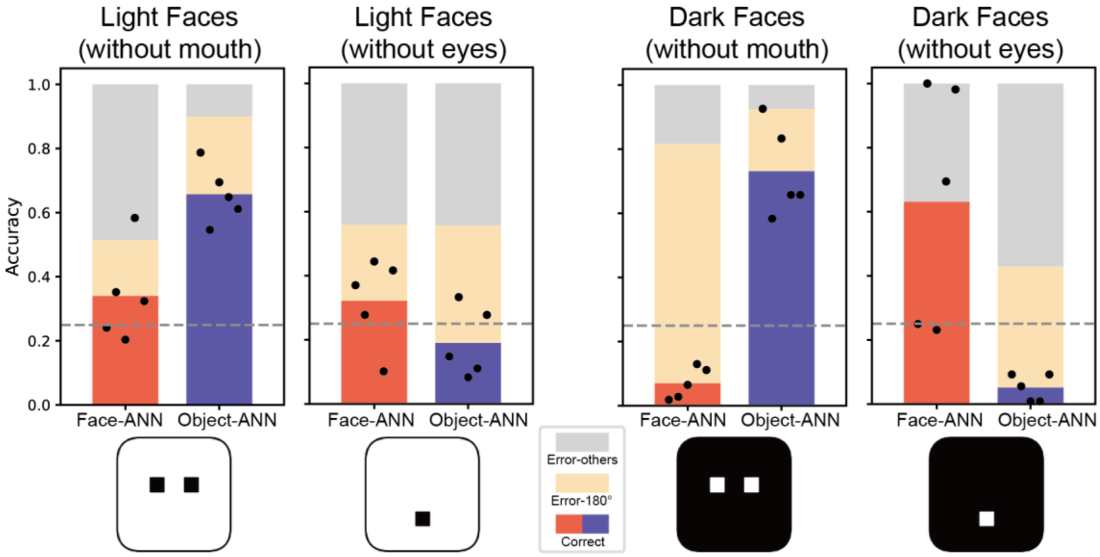


**Fig. S1. Model performance on degraded Three-dot Patterns.** The Face-ANN shows poor and unstable performance across all degraded conditions. For the Object-ANN, removal of the eye dots disrupts performance, whereas removal of the mouth dot does not affect classification accuracy.

#### Validating the effect of dataset size differences

In the main training procedure, the Face-ANN and Object-ANN were trained on the VGGFace2 and ImageNet datasets, respectively, which differ in size (approximately 3 million images for VGGFace2 and 1.2 million images for ImageNet). To examine whether this difference in dataset size influenced our results, we constructed a subset of VGGFace2 by selecting the first 3,500 identities, yielding approximately 1.2 million images. Using the same training parameters as in the main text, we retrained two models with different random seeds on this subset. The table below shows the performance of these models. Despite minor random fluctuations, their performance matched that of the Face-ANN reported in the main text and remained high on tasks where the Face-ANN originally excelled.

**Table S1. Performance comparison between size-matched VGGFace2 subset models (two random seeds), the original Face-ANNs, and the Object-ANNs.**

| | **Face-ANN** | **Face-ANN-subset** | | **Object-ANN** |
|---|---|---|---|---|
| **Stimulus Type** | *mean* | *seed=42* | *seed=108* | *mean* |
| Standard Faces | **1.000** | 1.000 | 1.000 | **1.000** |
| Thatcherized Faces | **0.993** | 0.978 | 0.989 | **1.000** |
| Infant Faces | **0.983** | 1.000 | 0.982 | 0.570 |
| Mooney Faces | **0.803** | 0.792 | 0.797 | 0.260 |
| Mooney Controls | 0.443 | 0.443 | 0.461 | 0.173 |
| Light Faces | **0.804** | 0.709 | 0.780 | 0.584 |
| Dark Faces | 0.338 | 0.504 | 0.357 | 0.428 |
| Arcimboldo Paintings | 0.573 | 0.577 | 0.577 | **1.000** |
| Face-like (designed) | 0.220 | 0.256 | 0.279 | **1.000** |
| Face-like (accidental) | 0.302 | 0.302 | 0.270 | **1.000** |
| Frontal Car | 0.178 | 0.128 | 0.152 | **1.000** |
| Cebus apella | 0.870 | 0.844 | 0.819 | **1.000** |
| Macaca fascicularis | 0.844 | 0.907 | 0.903 | **1.000** |
| Dog | 0.622 | 0.655 | 0.602 | **1.000** |
| Cat | 0.413 | 0.333 | 0.334 | **1.000** |
| Wolf | 0.559 | 0.540 | 0.477 | **1.000** |
| Fox | 0.188 | 0.152 | 0.155 | **1.000** |
| Leopard | 0.408 | 0.322 | 0.378 | **1.000** |

### Grayscale image testing

We additionally evaluated model performance using grayscale versions of all stimulus categories. The table below compares the results obtained with RGB stimuli and their corresponding grayscale counterparts. Note that the RGB results reported here are identical to those presented in the main text; they are shown again in the form of confusion matrices for ease of comparison. For the Face-ANN, performances with grayscale images were generally slightly lower than with RGB images, and in some cases even showed marginal improvements. In contrast, the Object-ANN typically exhibited a larger performance drop under grayscale conditions, suggesting that its learned representations rely more strongly on color information across stimulus categories. Interestingly, for several stimulus types (Giuseppe Arcimboldo Paintings, Cebus apella, Macaca fascicularis, and Leopard), the Face-ANN performed worse than the Object-ANN with RGB images, but this pattern reversed for grayscale images, with the Face-ANN achieving higher accuracy.

**Table S2. Results comparison between grayscale and RGB images.** Confusion matrices are shown, with rows indicating stimulus orientation and columns indicating the predicted orientation; values along the diagonal therefore represent the number of correctly classified images. Values are reported as mean ± SD across five models.

| | Grayscale | | | | | | | | RGB | | | | | | | |
|---|---|---|---|---|---|---|---|---|---|---|---|---|---|---|---|---|
| | Face-ANN | | | | Object-ANN | | | | Face-ANN | | | | Object-ANN | | | |
| | *be classified as* | | | | *be classified as* | | | | *be classified as* | | | | *be classified as* | | | |
| | *0°* | *90°* | *180°* | *270°* | *0°* | *90°* | *180°* | *270°* | *0°* | *90°* | *180°* | *270°* | *0°* | *90°* | *180°* | *270°* |
| **Standard Faces** | | | | | | | | | | | | | | | | |
| *stimuli* *0°* | 159.0±0.0 | 0.0±0.0 | 0.0±0.0 | 0.0±0.0 | 134.0±6.3 | 1.0±1.5 | 23.8±5.4 | 0.2±0.4 | 159.0±0.0 | 0.0±0.0 | 0.0±0.0 | 0.0±0.0 | 159.0±0.0 | 0.0±0.0 | 0.0±0.0 | 0.0±0.0 |
| *90°* | 0.0±0.0 | 159.0±0.0 | 0.0±0.0 | 0.0±0.0 | 0.4±0.8 | 138.0±10.2 | 1.6±1.9 | 19.0±9.3 | 0.0±0.0 | 159.0±0.0 | 0.0±0.0 | 0.0±0.0 | 0.0±0.0 | 159.0±0.0 | 0.0±0.0 | 0.0±0.0 |
| *180°* | 0.0±0.0 | 0.0±0.0 | 159.0±0.0 | 0.0±0.0 | 15.6±7.5 | 0.6±1.2 | 142.2±8.4 | 0.6±0.8 | 0.0±0.0 | 0.0±0.0 | 159.0±0.0 | 0.0±0.0 | 0.0±0.0 | 0.0±0.0 | 159.0±0.0 | 0.0±0.0 |
| *270°* | 0.0±0.0 | 0.0±0.0 | 0.0±0.0 | 159.0±0.0 | 0.6±0.8 | 16.6±7.3 | 1.2±1.9 | 140.6±7.8 | 0.0±0.0 | 0.0±0.0 | 0.0±0.0 | 159.0±0.0 | 0.0±0.0 | 0.0±0.0 | 0.0±0.0 | 159.0±0.0 |
| **Thatcherized Faces** | | | | | | | | | | | | | | | | |
| *stimuli* *0°* | 152.8±3.6 | 0.0±0.0 | 6.2±3.6 | 0.0±0.0 | 116.2±10.0 | 0.8±1.2 | 41.8±8.9 | 0.2±0.4 | 157.8±1.0 | 0.0±0.0 | 1.2±1.0 | 0.0±0.0 | 159.0±0.0 | 0.0±0.0 | 0.0±0.0 | 0.0±0.0 |
| *90°* | 0.0±0.0 | 152.8±3.3 | 0.0±0.0 | 6.2±3.3 | 0.4±0.8 | 124.6±20.6 | 0.4±0.5 | 33.6±20.2 | 0.0±0.0 | 157.0±1.7 | 0.0±0.0 | 2.0±1.7 | 0.0±0.0 | 159.0±0.0 | 0.0±0.0 | 0.0±0.0 |
| *180°* | 4.8±3.8 | 0.0±0.0 | 154.2±3.8 | 0.0±0.0 | 29.8±14.7 | 0.4±0.8 | 128.4±14.9 | 0.4±0.5 | 0.4±0.5 | 0.0±0.0 | 158.6±0.5 | 0.0±0.0 | 0.0±0.0 | 0.0±0.0 | 159.0±0.0 | 0.0±0.0 |
| *270°* | 0.0±0.0 | 4.6±3.9 | 0.0±0.0 | 154.4±3.9 | 0.4±0.5 | 29.2±17.6 | 0.8±1.6 | 128.6±18.1 | 0.0±0.0 | 0.8±1.6 | 0.0±0.0 | 158.2±1.6 | 0.0±0.0 | 0.0±0.0 | 0.0±0.0 | 159.0±0.0 |
| **Infant Faces** | | | | | | | | | | | | | | | | |
| *stimuli* *0°* | 104.6±1.4 | 0.0±0.0 | 4.4±1.4 | 0.0±0.0 | 44.0±3.3 | 21.0±1.3 | 24.8±5.1 | 19.2±3.0 | 107.2±1.2 | 0.2±0.4 | 1.6±1.4 | 0.0±0.0 | 69.6±9.3 | 16.6±4.5 | 16.4±4.3 | 6.4±2.7 |
| *90°* | 0.0±0.0 | 104.8±2.0 | 0.0±0.0 | 4.2±2.0 | 18.6±3.9 | 43.0±4.4 | 22.0±1.9 | 25.4±6.0 | 0.0±0.0 | 107.4±1.0 | 0.0±0.0 | 1.6±1.0 | 14.8±4.8 | 62.0±6.7 | 19.4±5.0 | 12.8±2.5 |
| *180°* | 3.8±1.5 | 0.0±0.0 | 105.0±1.3 | 0.2±0.4 | 21.8±3.4 | 20.8±2.9 | 46.8±8.5 | 19.6±2.6 | 1.6±1.4 | 0.0±0.0 | 107.2±1.2 | 0.2±0.4 | 23.8±6.6 | 13.4±4.7 | 60.4±1.0 | 11.4±3.4 |
| *270°* | 0.0±0.0 | 4.6±1.0 | 0.0±0.0 | 104.4±1.0 | 18.8±3.8 | 20.6±3.8 | 22.0±2.5 | 47.6±4.0 | 0.0±0.0 | 2.0±0.9 | 0.0±0.0 | 107.0±0.9 | 21.0±7.0 | 18.8±7.5 | 12.8±3.5 | 56.4±4.6 |

(Continued table)

| | | Grayscale | | | | | | | | RGB | | | | | | | |
|---|---|---|---|---|---|---|---|---|---|---|---|---|---|---|---|---|---|
| | | Face-ANN | | | | Object-ANN | | | | Face-ANN | | | | Object-ANN | | | |
| | | *be classified as* | | | | *be classified as* | | | | *be classified as* | | | | *be classified as* | | | |
| | | *0°* | *90°* | *180°* | *270°* | *0°* | *90°* | *180°* | *270°* | *0°* | *90°* | *180°* | *270°* | *0°* | *90°* | *180°* | *270°* |
| **Giuseppe Arcimboldo Paintings** | | | | | | | | | | | | | | | | | |
| *stimuli* | *0°* | 7.2±0.4 | 3.2±1.0 | 2.2±0.4 | 0.4±0.5 | 4.0±1.4 | 3.8±0.7 | 3.6±0.8 | 1.6±0.5 | 7.2±0.4 | 3.4±0.5 | 2.2±0.4 | 0.2±0.4 | 13.0±0.0 | 0.0±0.0 | 0.0±0.0 | 0.0±0.0 |
| | *90°* | 0.4±0.8 | 7.6±0.5 | 2.6±1.0 | 2.4±0.5 | 2.2±0.7 | 3.6±1.0 | 3.8±0.4 | 3.4±0.5 | 0.4±0.5 | 7.4±0.5 | 2.8±0.4 | 2.4±0.5 | 0.0±0.0 | 13.0±0.0 | 0.0±0.0 | 0.0±0.0 |
| | *180°* | 2.0±0.0 | 0.2±0.4 | 7.2±0.7 | 3.6±0.8 | 3.0±0.0 | 2.4±0.5 | 4.6±0.5 | 3.0±0.6 | 2.2±0.4 | 0.2±0.4 | 7.4±0.5 | 3.2±0.7 | 0.0±0.0 | 0.0±0.0 | 13.0±0.0 | 0.0±0.0 |
| | *270°* | 2.8±0.7 | 2.2±0.4 | 0.4±0.5 | 7.6±0.5 | 2.8±1.2 | 3.8±0.7 | 2.2±1.0 | 4.2±1.2 | 2.8±1.2 | 2.2±0.4 | 0.2±0.4 | 7.8±1.0 | 0.0±0.0 | 0.0±0.0 | 0.0±0.0 | 13.0±0.0 |
| **Face-like Objects (designed)** | | | | | | | | | | | | | | | | | |
| *stimuli* | *0°* | 21.4±1.0 | 15.4±2.9 | 17.8±4.0 | 23.4±1.0 | 37.6±1.7 | 9.6±3.0 | 18.4±1.4 | 12.4±1.9 | 15.6±0.8 | 18.4±5.7 | 20.6±2.3 | 23.4±4.1 | 78.0±0.0 | 0.0±0.0 | 0.0±0.0 | 0.0±0.0 |
| | *90°* | 19.8±1.7 | 22.4±2.5 | 16.4±1.2 | 19.4±2.0 | 10.2±1.0 | 39.4±2.9 | 9.8±1.6 | 18.6±2.7 | 22.6±1.6 | 17.0±2.5 | 18.8±1.6 | 19.6±2.4 | 0.0±0.0 | 78.0±0.0 | 0.0±0.0 | 0.0±0.0 |
| | *180°* | 18.8±1.7 | 21.8±3.0 | 20.0±1.9 | 17.4±3.1 | 19.6±2.0 | 10.6±1.0 | 39.0±2.0 | 8.8±1.7 | 19.4±2.0 | 23.2±3.4 | 18.0±1.9 | 17.4±2.5 | 0.0±0.0 | 0.0±0.0 | 78.0±0.0 | 0.0±0.0 |
| | *270°* | 18.8±1.3 | 18.6±1.5 | 18.2±1.2 | 22.4±1.4 | 9.0±1.7 | 18.8±2.5 | 11.6±2.1 | 38.6±1.6 | 17.8±4.3 | 20.0±2.5 | 22.2±2.5 | 18.0±1.9 | 0.0±0.0 | 0.0±0.0 | 0.0±0.0 | 78.0±0.0 |
| **Face-like Objects (accidental)** | | | | | | | | | | | | | | | | | |
| *stimuli* | *0°* | 23.4±1.2 | 18.6±2.2 | 22.2±1.2 | 21.8±2.0 | 45.0±2.3 | 11.0±1.3 | 22.4±1.9 | 7.6±0.5 | 24.2±3.0 | 17.2±1.9 | 23.6±2.6 | 21.0±2.2 | 86.0±0.0 | 0.0±0.0 | 0.0±0.0 | 0.0±0.0 |
| | *90°* | 21.6±1.4 | 23.0±1.9 | 18.0±1.8 | 23.4±1.9 | 6.0±0.9 | 46.6±1.4 | 8.8±1.2 | 24.6±1.0 | 19.6±1.5 | 26.4±2.2 | 17.4±2.6 | 22.6±1.9 | 0.0±0.0 | 86.0±0.0 | 0.0±0.0 | 0.0±0.0 |
| | *180°* | 24.6±2.1 | 19.6±3.9 | 24.0±3.6 | 17.8±2.2 | 22.6±3.5 | 7.8±1.2 | 47.0±2.6 | 8.6±0.8 | 22.6±3.3 | 20.8±4.3 | 27.0±1.3 | 15.6±2.0 | 0.0±0.0 | 0.0±0.0 | 86.0±0.0 | 0.0±0.0 |
| | *270°* | 17.4±0.5 | 23.6±2.1 | 22.2±1.8 | 22.8±3.5 | 8.4±1.4 | 25.4±1.6 | 6.8±1.6 | 45.4±2.2 | 15.0±1.4 | 22.2±2.5 | 22.6±1.5 | 26.2±2.7 | 0.0±0.0 | 0.0±0.0 | 0.0±0.0 | 86.0±0.0 |
| **Frontal Car** | | | | | | | | | | | | | | | | | |
| *stimuli* | *0°* | 21.8±7.5 | 21.6±9.0 | 100.6±7.6 | 16.0±4.9 | 137.0±2.8 | 2.8±1.9 | 14.2±1.9 | 6.0±1.8 | 27.4±7.1 | 20.6±10.3 | 97.8±9.5 | 14.2±5.8 | 159.8±0.4 | 0.0±0.0 | 0.0±0.0 | 0.2±0.4 |
| | *90°* | 14.6±2.2 | 23.0±6.4 | 20.2±4.3 | 102.2±6.0 | 3.6±1.2 | 136.8±4.5 | 4.2±1.3 | 15.4±3.5 | 12.6±1.9 | 29.6±6.5 | 20.8±6.5 | 97.0±10.0 | 0.0±0.0 | 160.0±0.0 | 0.0±0.0 | 0.0±0.0 |
| | *180°* | 96.8±9.7 | 16.0±5.9 | 25.2±3.3 | 22.0±3.7 | 15.4±4.5 | 3.6±1.0 | 136.4±5.7 | 4.6±1.5 | 94.6±12.4 | 15.6±5.3 | 30.2±3.3 | 19.6±5.0 | 0.0±0.0 | 0.0±0.0 | 160.0±0.0 | 0.0±0.0 |
| | *270°* | 22.0±5.8 | 97.6±9.4 | 15.0±2.8 | 25.4±2.9 | 2.8±1.9 | 16.0±1.7 | 4.6±1.4 | 136.6±3.4 | 20.2±7.3 | 99.0±13.3 | 13.8±4.3 | 27.0±3.7 | 0.0±0.0 | 0.0±0.0 | 0.0±0.0 | 160.0±0.0 |
| **Cebus apella** | | | | | | | | | | | | | | | | | |
| *stimuli* | *0°* | 138.2±5.2 | 3.8±1.6 | 14.8±2.9 | 5.2±1.9 | 98.0±10.1 | 25.8±6.5 | 15.4±2.6 | 22.8±3.1 | 139.8±4.0 | 4.6±1.0 | 12.0±2.3 | 5.6±2.1 | 162.0±0.0 | 0.0±0.0 | 0.0±0.0 | 0.0±0.0 |
| | *90°* | 5.6±1.0 | 137.6±4.6 | 3.8±0.7 | 15.0±4.3 | 24.8±3.4 | 92.8±8.9 | 27.4±2.7 | 17.0±5.2 | 5.4±0.5 | 138.6±3.7 | 4.8±0.7 | 13.2±3.5 | 0.0±0.0 | 162.0±0.0 | 0.0±0.0 | 0.0±0.0 |
| | *180°* | 14.4±3.8 | 3.8±1.7 | 141.0±4.5 | 2.8±0.4 | 16.4±2.0 | 23.8±2.3 | 97.6±3.2 | 24.2±3.2 | 12.6±6.4 | 3.6±0.5 | 141.8±7.0 | 4.0±0.9 | 0.0±0.0 | 0.0±0.0 | 162.0±0.0 | 0.0±0.0 |
| | *270°* | 2.4±1.5 | 12.0±3.1 | 5.0±1.4 | 142.6±5.4 | 26.8±4.9 | 16.2±5.2 | 23.8±2.6 | 95.2±7.8 | 3.4±1.9 | 11.0±2.6 | 4.2±0.7 | 143.4±3.7 | 0.0±0.0 | 0.0±0.0 | 0.0±0.0 | 162.0±0.0 |
| **Macaca fascicularis** | | | | | | | | | | | | | | | | | |
| *stimuli* | *0°* | 60.2±2.7 | 1.0±0.6 | 11.8±4.4 | 2.0±1.3 | 52.8±1.7 | 9.0±3.0 | 6.2±2.1 | 7.0±1.7 | 63.8±1.5 | 1.0±0.9 | 9.0±2.9 | 1.2±0.7 | 75.0±0.0 | 0.0±0.0 | 0.0±0.0 | 0.0±0.0 |
| | *90°* | 2.0±1.1 | 60.6±3.6 | 1.4±0.5 | 11.0±4.7 | 7.8±2.7 | 53.6±4.1 | 8.4±2.2 | 5.2±1.6 | 1.4±1.0 | 63.4±2.2 | 1.2±0.7 | 9.0±2.9 | 0.0±0.0 | 75.0±0.0 | 0.0±0.0 | 0.0±0.0 |
| | *180°* | 10.8±2.7 | 1.6±1.0 | 61.4±2.9 | 1.2±0.4 | 5.4±2.9 | 7.2±2.9 | 56.6±5.8 | 5.8±1.2 | 9.6±2.9 | 1.4±1.0 | 62.8±2.7 | 1.2±0.4 | 0.0±0.0 | 0.0±0.0 | 75.0±0.0 | 0.0±0.0 |
| | *270°* | 1.2±0.4 | 12.0±2.1 | 2.2±1.3 | 59.6±1.5 | 9.4±1.0 | 6.4±2.1 | 8.0±1.7 | 51.2±3.1 | 1.0±0.6 | 9.0±1.7 | 1.8±1.3 | 63.2±1.2 | 0.0±0.0 | 0.0±0.0 | 0.0±0.0 | 75.0±0.0 |

(Continued table)

| | Grayscale | | | | | | | | RGB | | | | | | | |
|---|---|---|---|---|---|---|---|---|---|---|---|---|---|---|---|---|
| | Face-ANN | | | | Object-ANN | | | | Face-ANN | | | | Object-ANN | | | |
| | *be classified as* | | | | *be classified as* | | | | *be classified as* | | | | *be classified as* | | | |
| | *0°* | *90°* | *180°* | *270°* | *0°* | *90°* | *180°* | *270°* | *0°* | *90°* | *180°* | *270°* | *0°* | *90°* | *180°* | *270°* |
| **Dog** | | | | | | | | | | | | | | | | |
| *stimuli 0°* | 102.0±2.3 | 6.6±1.2 | 46.2±2.8 | 5.2±0.7 | 125.8±3.3 | 9.4±2.6 | 18.2±1.5 | 6.6±2.0 | 100.4±1.6 | 10.0±1.5 | 44.6±1.0 | 5.0±0.0 | 160.0±0.0 | 0.0±0.0 | 0.0±0.0 | 0.0±0.0 |
| *90°* | 3.8±0.7 | 100.6±7.6 | 6.2±1.7 | 49.4±7.7 | 6.4±1.9 | 127.2±2.3 | 9.8±0.7 | 16.6±2.4 | 5.4±1.2 | 98.4±2.9 | 8.0±1.5 | 48.2±3.1 | 0.0±0.0 | 160.0±0.0 | 0.0±0.0 | 0.0±0.0 |
| *180°* | 47.8±4.7 | 4.2±0.7 | 101.2±6.0 | 6.8±1.7 | 16.6±2.6 | 5.8±0.4 | 129.0±3.2 | 8.6±3.1 | 46.4±1.4 | 6.0±2.6 | 97.2±3.2 | 10.4±1.7 | 0.0±0.0 | 0.0±0.0 | 160.0±0.0 | 0.0±0.0 |
| *270°* | 6.2±1.6 | 45.0±2.6 | 4.8±1.0 | 104.0±2.8 | 8.4±1.5 | 16.2±2.0 | 6.0±1.9 | 129.4±3.9 | 7.8±1.0 | 45.0±3.8 | 5.4±1.2 | 101.8±4.4 | 0.0±0.0 | 0.0±0.0 | 0.0±0.0 | 160.0±0.0 |
| **Cat** | | | | | | | | | | | | | | | | |
| *stimuli 0°* | 52.2±8.7 | 27.6±3.6 | 52.8±11.1 | 27.4±1.4 | 100.6±4.9 | 18.4±5.0 | 26.6±4.0 | 14.4±1.2 | 65.4±7.9 | 30.0±2.4 | 40.4±8.2 | 24.2±1.8 | 160.0±0.0 | 0.0±0.0 | 0.0±0.0 | 0.0±0.0 |
| *90°* | 23.0±3.8 | 47.8±8.1 | 30.4±5.2 | 58.8±13.3 | 17.2±3.4 | 95.4±5.5 | 20.6±2.4 | 26.8±1.7 | 22.4±2.6 | 63.6±6.9 | 28.6±4.1 | 45.4±9.0 | 0.0±0.0 | 160.0±0.0 | 0.0±0.0 | 0.0±0.0 |
| *180°* | 53.8±10.9 | 25.0±5.7 | 48.8±10.6 | 32.4±4.2 | 25.0±2.4 | 14.6±0.5 | 100.0±8.7 | 20.4±7.6 | 43.6±5.5 | 21.2±5.7 | 65.4±6.2 | 29.8±2.2 | 0.0±0.0 | 0.0±0.0 | 160.0±0.0 | 0.0±0.0 |
| *270°* | 28.6±2.0 | 53.4±14.9 | 25.2±2.9 | 52.8±13.3 | 18.4±3.5 | 23.2±1.3 | 15.6±2.4 | 102.8±1.3 | 25.8±2.7 | 41.2±8.6 | 23.4±0.5 | 69.6±9.7 | 0.0±0.0 | 0.0±0.0 | 0.0±0.0 | 160.0±0.0 |
| **Wolf** | | | | | | | | | | | | | | | | |
| *stimuli 0°* | 38.6±2.4 | 1.4±1.0 | 34.4±1.9 | 0.6±0.5 | 55.2±2.4 | 7.2±1.6 | 4.0±1.4 | 8.6±1.6 | 41.2±3.4 | 2.2±1.7 | 30.6±3.3 | 1.0±0.9 | 75.0±0.0 | 0.0±0.0 | 0.0±0.0 | 0.0±0.0 |
| *90°* | 0.4±0.5 | 39.8±1.5 | 1.0±0.6 | 33.8±1.3 | 9.8±1.7 | 54.0±3.2 | 6.2±1.3 | 5.0±1.1 | 0.8±0.7 | 42.8±2.4 | 0.8±0.7 | 30.6±2.2 | 0.0±0.0 | 75.0±0.0 | 0.0±0.0 | 0.0±0.0 |
| *180°* | 35.0±3.2 | 0.4±0.5 | 37.8±2.9 | 1.8±1.6 | 4.0±0.6 | 10.0±1.7 | 53.8±4.3 | 7.2±2.8 | 32.4±3.7 | 0.4±0.5 | 40.6±3.1 | 1.6±1.5 | 0.0±0.0 | 0.0±0.0 | 75.0±0.0 | 0.0±0.0 |
| *270°* | 1.6±1.4 | 32.8±3.1 | 0.6±0.8 | 40.0±1.8 | 6.4±2.7 | 5.6±1.0 | 10.2±1.0 | 52.8±2.5 | 1.2±0.7 | 30.0±3.4 | 0.6±0.8 | 43.2±2.6 | 0.0±0.0 | 0.0±0.0 | 0.0±0.0 | 75.0±0.0 |
| **Fox** | | | | | | | | | | | | | | | | |
| *stimuli 0°* | 29.0±5.2 | 1.8±0.4 | 126.4±4.5 | 2.8±1.2 | 84.6±3.9 | 19.6±1.6 | 34.0±4.9 | 21.8±1.6 | 31.4±7.1 | 2.2±0.4 | 122.8±7.0 | 3.6±0.8 | 160.0±0.0 | 0.0±0.0 | 0.0±0.0 | 0.0±0.0 |
| *90°* | 3.2±1.3 | 30.4±1.9 | 1.6±1.0 | 124.8±3.7 | 24.8±3.5 | 82.2±5.4 | 19.2±2.8 | 33.8±2.7 | 4.4±1.0 | 31.2±4.4 | 3.4±1.9 | 121.0±4.6 | 0.0±0.0 | 160.0±0.0 | 0.0±0.0 | 0.0±0.0 |
| *180°* | 129.2±4.2 | 4.4±0.8 | 24.8±3.2 | 1.6±1.2 | 36.0±1.7 | 23.2±1.6 | 84.4±4.2 | 16.4±3.0 | 122.0±5.4 | 5.8±1.7 | 28.8±3.9 | 3.4±2.3 | 0.0±0.0 | 0.0±0.0 | 160.0±0.0 | 0.0±0.0 |
| *270°* | 1.2±0.7 | 129.2±2.7 | 2.8±1.6 | 26.8±4.1 | 19.2±1.3 | 36.0±1.4 | 22.6±2.2 | 82.2±3.4 | 2.6±1.0 | 124.6±5.7 | 4.2±2.0 | 28.6±6.9 | 0.0±0.0 | 0.0±0.0 | 0.0±0.0 | 160.0±0.0 |
| **Leopard** | | | | | | | | | | | | | | | | |
| *stimuli 0°* | 77.6±3.9 | 37.6±3.6 | 8.8±2.7 | 36.0±1.5 | 48.8±2.1 | 9.4±4.5 | 86.4±4.0 | 15.4±3.7 | 62.2±4.1 | 8.0±2.6 | 73.6±4.1 | 16.2±3.0 | 160.0±0.0 | 0.0±0.0 | 0.0±0.0 | 0.0±0.0 |
| *90°* | 35.0±5.2 | 81.0±6.4 | 36.4±7.1 | 7.6±2.3 | 14.2±5.5 | 60.2±12.8 | 10.0±4.2 | 75.6±7.2 | 14.2±4.5 | 69.0±12.8 | 7.6±2.6 | 69.2±7.1 | 0.0±0.0 | 160.0±0.0 | 0.0±0.0 | 0.0±0.0 |
| *180°* | 7.6±3.4 | 31.8±3.7 | 84.4±10.1 | 36.2±5.6 | 76.4±12.6 | 15.4±4.1 | 57.6±12.4 | 10.6±2.9 | 68.8±9.0 | 14.2±5.5 | 66.2±10.3 | 10.8±3.5 | 0.0±0.0 | 0.0±0.0 | 160.0±0.0 | 0.0±0.0 |
| *270°* | 34.8±4.4 | 6.8±1.6 | 35.4±5.7 | 83.0±2.5 | 8.6±2.9 | 80.4±10.1 | 15.0±4.3 | 56.0±11.9 | 7.4±2.6 | 74.6±11.4 | 14.2±2.6 | 63.8±12.1 | 0.0±0.0 | 0.0±0.0 | 0.0±0.0 | 160.0±0.0 |
